# A mutation at the dimer interface regulates enzyme catalysis

**DOI:** 10.1101/2025.04.29.650001

**Authors:** Caitlin E Hatton, Laura Falkenburg, Pedram Mehrabi

## Abstract

Fluoroacetate Dehalogenase (FAcD), a homodimeric dehalogenase of interest in bioremediation, displays half-of-the-sites reactivity, requiring allosteric communication between the subunits to coordinate the reaction. Dimer interfaces are an obvious site of interest for understanding communication between subunits and have been shown, in a variety of enzyme systems, to mediate such communication. However, mutations at these interfaces often need to be substantial to noticeably affect protein activity. In this study, we demonstrate that two subtle interface mutations of FAcD, either accelerate the reaction or results in near-complete loss of activity with structural evidence of disruption at the dimer interface. By examining these variants using ultrahigh-resolution crystallography and kinetics studies, the influence of dimer interface variations in hydrogen-bonding networks on enzyme activity can be elucidated.

## Introduction

Fluoroacetate dehalogenase (FAcD) is a homodimeric protein from *Rhodopseudomonas palustris* that catalyses the conversation of toxic halogenated-acetates into glycolate and acid, particularly fluoroacetate (FAc), by breaking the strongest single bond in organic chemistry (C-F) (Figure 1a)^1^. FAcD catalyses this reaction through an SN_2_ involving the conserved catalytic triad of Asn110, Asn134, His280 (Figure 1a). Understanding how the enzyme performs this reaction under mild conditions is important for bioremediation and the removal of toxic compounds from the environment. The oligomeric state of FAcD plays a crucial role in modulating its enzymatic activity.

**Figure 1.**
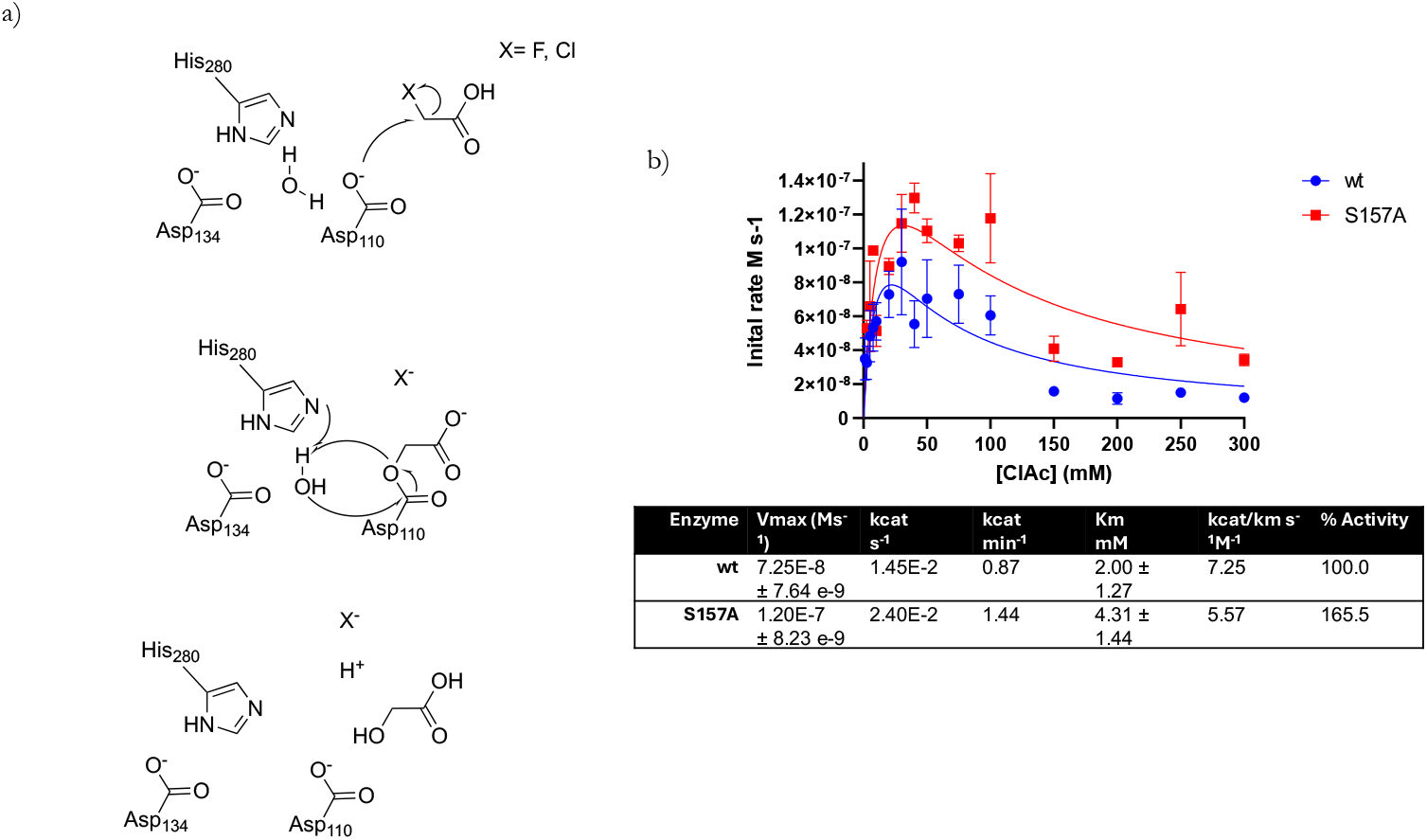
Reaction and kinetics of FAcD. a) Fluoroacetate and chloroacetate are converted into glycolate, and FCl or HCl respectively. Catalysis involves a conserved catalytic triad of Asp110, Asp134, His280 and occurs via a S_N_2 reaction. b) Enzyme kinetics of FAcD with chloroacetate, monitoring the production of H^+^. Initial rates show that FAcD exhibits substrate inhibition at higher [ClAc], the Ser157Ala (red) manages to achieve a higher V_max_ compared to WT (blue).

Oligomerization is a fundamental feature of many proteins, with dimers being among the most common forms ^2,3^. It influences various aspects of enzyme function, including substrate binding, entropy compensation, catalytic efficiency, stability, and allosteric regulation. Oligomeric enzymes often exhibit allosteric behaviour, where long-distance interactions control enzyme function through events such as ligand binding. This allows allosteric enzymes to rapidly regulate protein activity as a response to cellular needs.

Mutations at subunit interfaces have been shown to alter enzyme activity in oligomeric systems, often through changes in cooperativity, stability, or conformational heterogeneity. For example, introducing a tryptophan at the dimer interface in glutathione reductase (GR) induces positive cooperativity between the glutathione binding sites of each subunit, at the expense of thermal stability, without disrupting oligomerisation ^4^. Similarly, mutations at the dimer interface of Human immunodeficiency virus type 1 (HIV-1) protease have been shown to affect both catalytic activity and turnover, although these changes are also associated with decreased dimer stability ^5^. In glyceraldehyde 3-phosophate dehydrogenase (GAPDH), a dimer interface mutation that mimics a post-translational modification disrupts the tetramer’s ability to bind RNA, likely by preventing higher-order oligomerisation ^6^. Likewise, mutations at the interface between domains of pyruvate kinase (PK) not only alter enzymatic activity but also induce structural rearrangements ^7^.

The FAcD homodimer exhibits substrate inhibition at high substrate concentrations and displays half-of-the-sites reactivity, a form of negative cooperativity, where only one monomer is active at a time. Previous work on FAcD has shown that FAcD binds substrate in one active site to initiate the reaction. Then only when the covalent intermediate has formed in the first active site does the second active site begin to populate with substrate and begin turnover. This is before the first active site has released product ^8^. This mechanism allows the bacterium to tightly regulate the balance between harnessing halogenated acetate as a carbon source with the pH drop due to the acid product. FAcD is, unsurprisingly, an obligate homodimer. Simultaneous mutation of a large number of residues at the dimer interface is required to yield a functional monomeric form of FAcD ^9^. The dimeric interface of FAcD therefore presents an ideal opportunity for investigating allosteric communication.

It has previously been shown that asymmetry between the FAcD monomers drives catalysis and facilitates communication between the subunits ^3,8^. Specifically, rotation of a serine side chain at the dimer interface (Ser157) correlates with the progression of the reaction, accompanied by changes in water coordination at the interface ^8^. Ser157 was therefore predicted to play a key role in allosteric communication in FAcD, by transmitting information about the reaction state from one subunit to the other via an allosteric pathway. To gain further insight into this residue, Ser157 variants were generated and ultrahigh-resolution structures (<0.9 Å) determined, allowing better visualisation of hydrogens and identification of alternative conformations representing low-population conformers that are invisible at lower resolutions^10^. This level of detail provides a rare view into how specific structural features contribute to protein function. Notably, even conservative mutations of Ser157 led to significant changes in enzymatic activity, highlighting the impact of subtle variations in hydrogen-bond networks far from the active site on catalysis.

## Results

To investigate the role of Ser157 in FAcD catalysis, it was mutated to threonine and alanine. Colourimetric monitoring of proton production was used to measure the enzyme turnover of chloroacetate (ClAc) by both WT and mutant variants of FAcD (Figure 1b). As previously reported, WT FAcD exhibits substrate inhibition at high concentrations of ClAc. The Ser157Ala variant shows a higher turnover rate for ClAc and maintains WT-like substrate inhibition. In contrast, no turnover could be detected for the Ser157Thr variant, resembling the behaviour observed for Asp110Asn and His280Asn mutants. These residues are part of the conserved active site catalytic triad (Asp110, Asp134, His280) and their mutation results in enzyme inactivation (Table 1 and Figure 1b).

**Table 1.** Kinetics of FAcD and mutants of ClAc hydrolysis. The parameters were determined from at least triplicate measurements, and the standard error of the mean is shown. The % activity normalizes all *k*_*c*at_ to ClAc hydrolysis by WT.

|  | WT_Apo | S157A_Apo | S157T_Apo | S157T_Complex |
| --- | --- | --- | --- | --- |
| Wavelength (Å) | 0.465 | 0.689 | 0.529 | 0.689 |
| Resolution range (Å) | 28.48 - 0.86 (0.94 - 0.86) | 35.33 - 0.85 (0.93 - 0.85) | 40.42 - 0.87 (0.95 - 0.87) | 56.85-0.89 (0.99-0.89) |
| Space group | P 1 21 1 | P 1 21 1 | P 1 21 1 | P 1 21 1 |
| Unit Cell (a,b,c) (Å) | 41.94,<br>79.48,<br>85.10 | 41.87,<br>78.50,<br>85.15 | 41.89,<br>78.25,<br>84.91 | 41.70,<br>78.13,<br>84.95, |
| Unit cell ($\alpha,\beta,\gamma$) (°) | 90,<br>103.17,<br>90 | 90,<br>103.65,<br>90 | 90,<br>103.19,<br>90 | 90,<br>102.61,<br>90 |
| Total reflections | 7289960<br>(338693) | 6687228<br>(318384) | 6706508<br>(345819) | 5556219<br>(274877) |
| Unique reflections | 360718<br>(18036) | 341382<br>(17069) | 337935<br>(16897) | 280924<br>(14046) |
| Multiplicity | 20.2 (18.8) | 19.6 (18.7) | 19.8 (20.5) | 19.8 (19.6) |
| Completeness (Spherical) (%) | 79.6 (17.9) | 70.8 (13.5) | 76.8 (16.5) | 69.5 (12.4) |
| Mean I/sigma(I) | 27.3 (1.6) | 25 (1.9) | 21.1 (1.8) | 23.4 (1.8) |
| Wilson B-factor (Å<sup>2</sup>) | 9.07 | 8.33 | 8.07 | 8.88 |
| CC1/2 | 1 (0.689) | 0.999 (0.859) | 0.99 (0.731) | 1 (0.788) |
| R-work | 0.1068 | 0.107 | 0.1133 | 0.1169 |
| R-free | 0.1252 | 0.1263 | 0.1341 | 0.1324 |
| Number of non-hydrogen atoms | 7615 | 7511 | 8071 | 7340 |
| macromolecules | 6422 | 6350 | 6970 | 6300 |
| ligands | 3 | 1 | 1 | 8 |
| solvent | 1190 | 1160 | 1100 | 1032 |
| Protein residues | 591 | 597 | 596 | 587 |
| RMS (bonds) (Å) | 0.092 | 0.092 | 0.091 | 0.091 |
| RMS (angles) (°) | 2.06 | 2.24 | 1.86 | 2.02 |
| Ramachandran favored (%) | 97.26 | 96.8 | 97.63 | 97.54 |
| Ramachandran allowed (%) | 2.57 | 3.04 | 2.37 | 2.28 |
| Ramachandran outliers (%) | 0.17 | 0.17 | 0 | 0.18 |
| Rotamer outliers (%) | 2 | 0.93 | 1.69 | 2.66 |
| Clashscore | 12.02 | 14.91 | 5.03 | 10.29 |
| Average B-factor (Å<sup>2</sup>) | 15.63 | 13.03 | 13.74 | 14.33 |
| macromolecules | 12.46 | 9.62 | 10.81 | 11.44 |
| ligands | 11.84 | 8.31 | 8.91 | 38.18 |
| solvent | 32.77 | 31.74 | 32.31 | 31.81 |

To examine the structural basis for the altered activity upon Ser157 mutation, ultrahigh-resolution structures (< 0.9 Å; refinement statistics in Table 2.) of substrate-free FAcD-WT, Ser157Ala, and Ser157Thr were determined (Table 2). At this resolution, hydrogen positions can be accurately assigned, and hydrogen-bonding networks clearly identified. Both Ser157 mutants maintained the expected dimeric structure. In WT FAcD, five structural waters are observed at the dimer interface (Figure 2a). W1 and W5 are “symmetry mates”, related by a non-crystallographic two-fold, and are tightly coordinated by the protein backbone. W2 and W4 are also similarly related and are coordinated by W1/W5 and the FAcD backbone. W3 is uniquely positioned at the interface without a symmetric counterpart. The Ser157Ala variant preserved the WT water coordination at the dimer interface with minimal change in water position, occupancy (Figure 2b) and B-factor (Figure 2c) of these waters when compared to WT, despite the loss of a hydroxyl functional group at residue 157 in this variant. Larger changes were observed for the Ser157Thr variant and are discussed below.

**Table 2.** Data and refinement statistics. Statistics for the highest resolution shell are given in parentheses.

**Figure 2.**
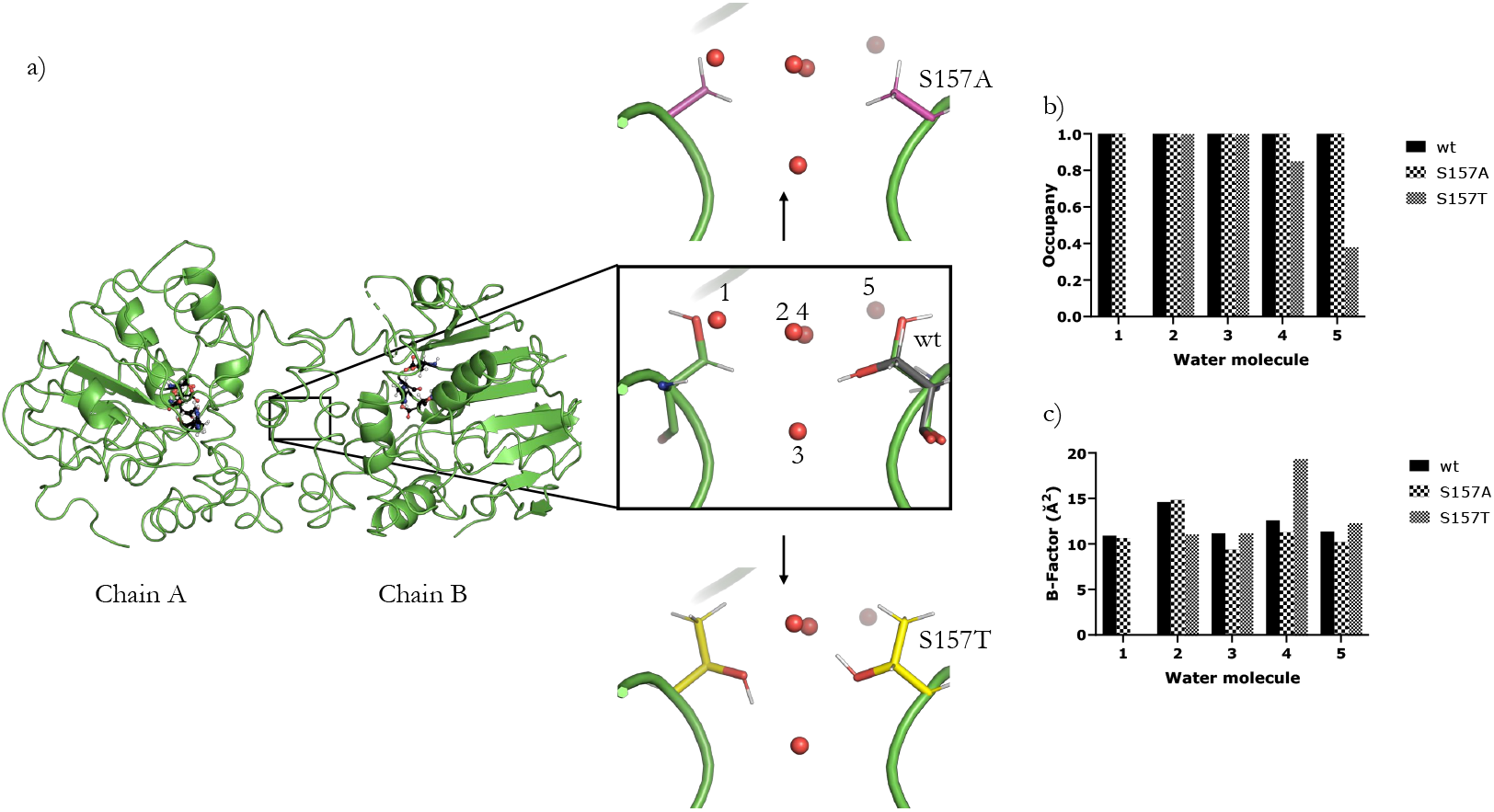
The FAcD dimer interface. a) WT (green), Ser157Ala (purple), and S157Thr (yellow). Water 1 is not present in Ser157Thr, and water 2 has been displaced and is found in an alternative conformation. b) Ser157Ala shows no change in occupancy compared to wt, whilst Ser157Thr shows that the occupancy of water 4 and 5 has reduced. c) B-factor of each water for WT, Ser157Ala, and Ser157Thr.

In WT FAcD, the Ser157 side chain in Chain A is adopts a single conformation, oriented towards Trp142. In Chain B the Ser157 side chain is found in two conformations, with the major conformation (0.78 occupancy) pointing in the same direction as in chain A, while the minor conformation (0.22 occupancy) is oriented toward Tyr154. In the Ser157Thr mutant, the hydroxyl moiety unambiguously points toward Tyr154 in both chain A and B (Figure 2a). Notably, W1 is absent in Ser157Thr, while its symmetry mate, W5 is present with significantly reduced occupancy (Figure 2b). The exclusion of W1 in Ser157Thr therefore cannot be due to a simple steric hindrance at the interface as this would also exclude its symmetry mate W5. In Ser157Thr W2 adopts two conformations and is displaced from its position in WT FAcD (Figure 2a), whereas its symmetry mate W4 remains unchanged. W3 also remains unchanged from WT.

In WT FAcD, W1 is coordinated in Chain A through backbone interactions with Ile153, Tyr154, Trp156, as well as side-chain interactions with Tyr141, Met145, and Ser157, (Figure 3a). Tyr141 in Chain A adopts two conformations, with the major conformation (0.65 occupancy) *π*-stacking with Trp156 and previously reported as the conformation when the subunit is primed to bind substrate. While mutation to Ser157Ala removes a hydrogen bond, due to the loss of the hydroxyl-moiety, all other interactions are retained, allowing W1 to remain coordinated (Figure 3b). While Tyr141 maintains *π*-stacking with Trp156 this conformation now has an occupancy of 0.77. In contrast, the Ser157Thr mutation abolishes coordination of W1 in Chain A. The combination of losing the Ser157 hydroxyl interaction and the associated rotation of Tyr141 appears sufficient to prevent stable W1 binding (Figure 3c). In this variant, Tyr141 no longer π-stacks with Trp156.

**Figure 3.**
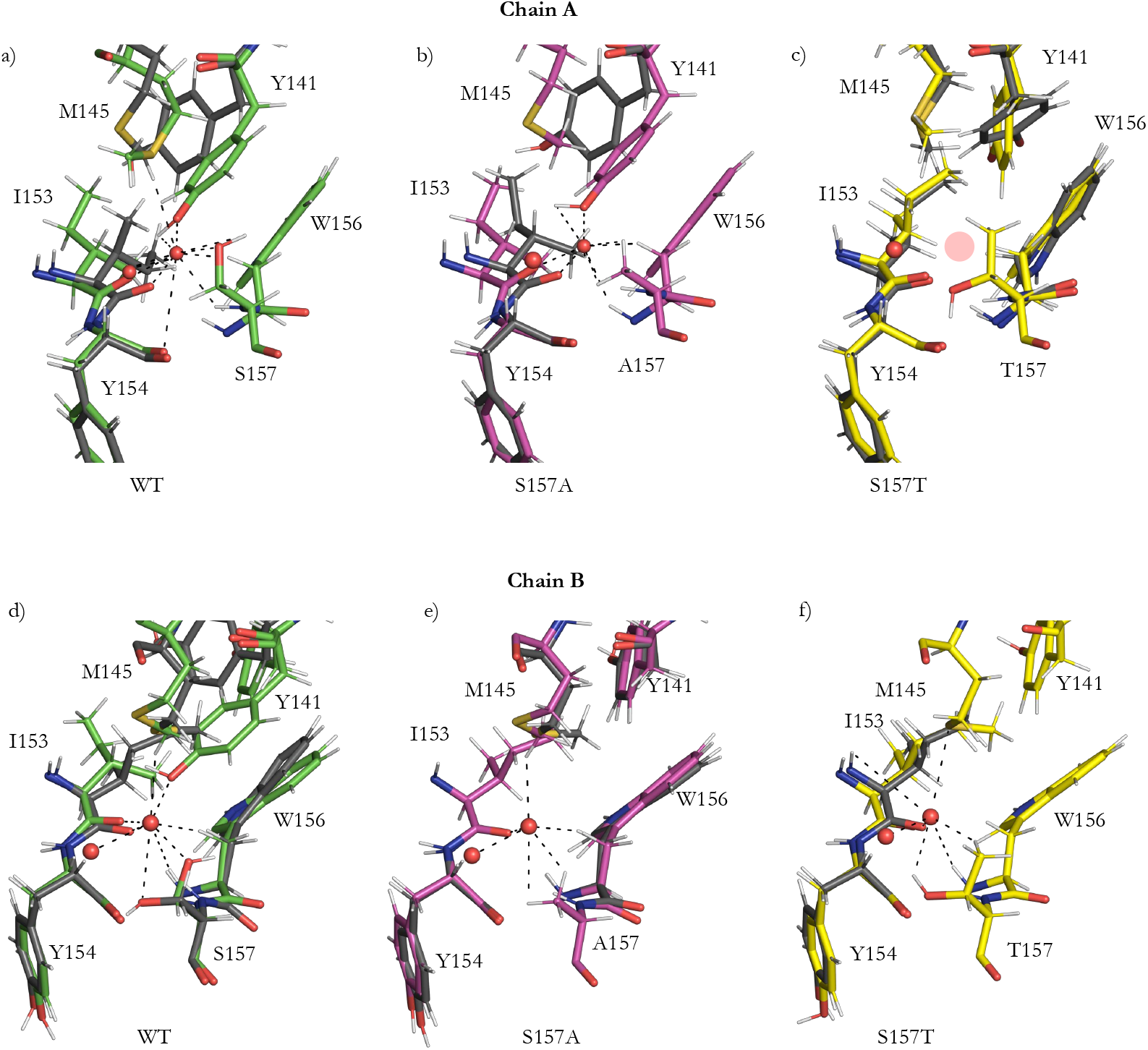
Water coordination for W1 and W5 at the FAcD dimer interface. Alternative conformations are depicted in grey. a)WT Chain A-W1 is coordinated by a variety of side and main chain interactions. b) Ser157Ala Chain A – The water coordination is largely unchanged compared to wt. c) Ser157Thr-Chain A - The water is not coordinated at all despite the interactions being maintained. d) WT Chain B - Water 5 is coordinated by a variety of side and main chain interactions. e) Ser157Ala Chain B – The water coordination is largely unchanged compared to WT. f) Ser157Thr Chain B-The water is still coordinated despite not being coordinated in Chain A.

In Chain B of WT, W5 (the equivalent to W1) is coordinated by the hydroxyl of Tyr141 (when in the *π*-stacking conformation at 0.22 occupancy), with additional interactions with the Met145 side chain, the backbone of Ile153, Ser157, and side chain of the major conformation of Ser157 (Figure 3d). In both Ser157Ala and Ser157Thr variants, Tyr141 is found only in a single conformation pointing away from the active site (Figure 3e, 3f). Closer inspection reveals that W5 in Ser157Thr does not interact with the hydroxyl of the Thr157 and Tyr141 side chains and lacks coordination from the Met145 side chain. Although Ser157Ala retains W5 coordination, these subtle differences in water coordination likely contribute to the reduction of water coordination at the interface of Ser157Thr (Figure 3c).

Closer analysis of the region surrounding the mutation site (residues 145-160) reveals that WT adopts two predominant conformations in approximately a 2:1 occupancy ratio. The Ser157Ala variant appears to favour a single conformation that aligns with the higher-occupancy state observed in the WT. In contrast, Ser157Thr variant also adopts two conformations, but neither overlap with those found in the WT or Ser157Ala. This observation is particularly evident with the Met145 and Tyr141 side chains, Tyr141 has been previously an indicator of conformational selectivity, and its conformational identifies the catalytic state. The side chain of Met145 in Ser157Thr is completely different to both WT and Ser157Ala in both chains (Figure 4a, 4b, 4c). The difference in conformation may explain the loss of interactions of these side chains, thereby changing water coordination at the interface. These subtle conformational differences are only discernible at ultrahigh resolution and would not be visible using other structural methods.

**Figure 4.**
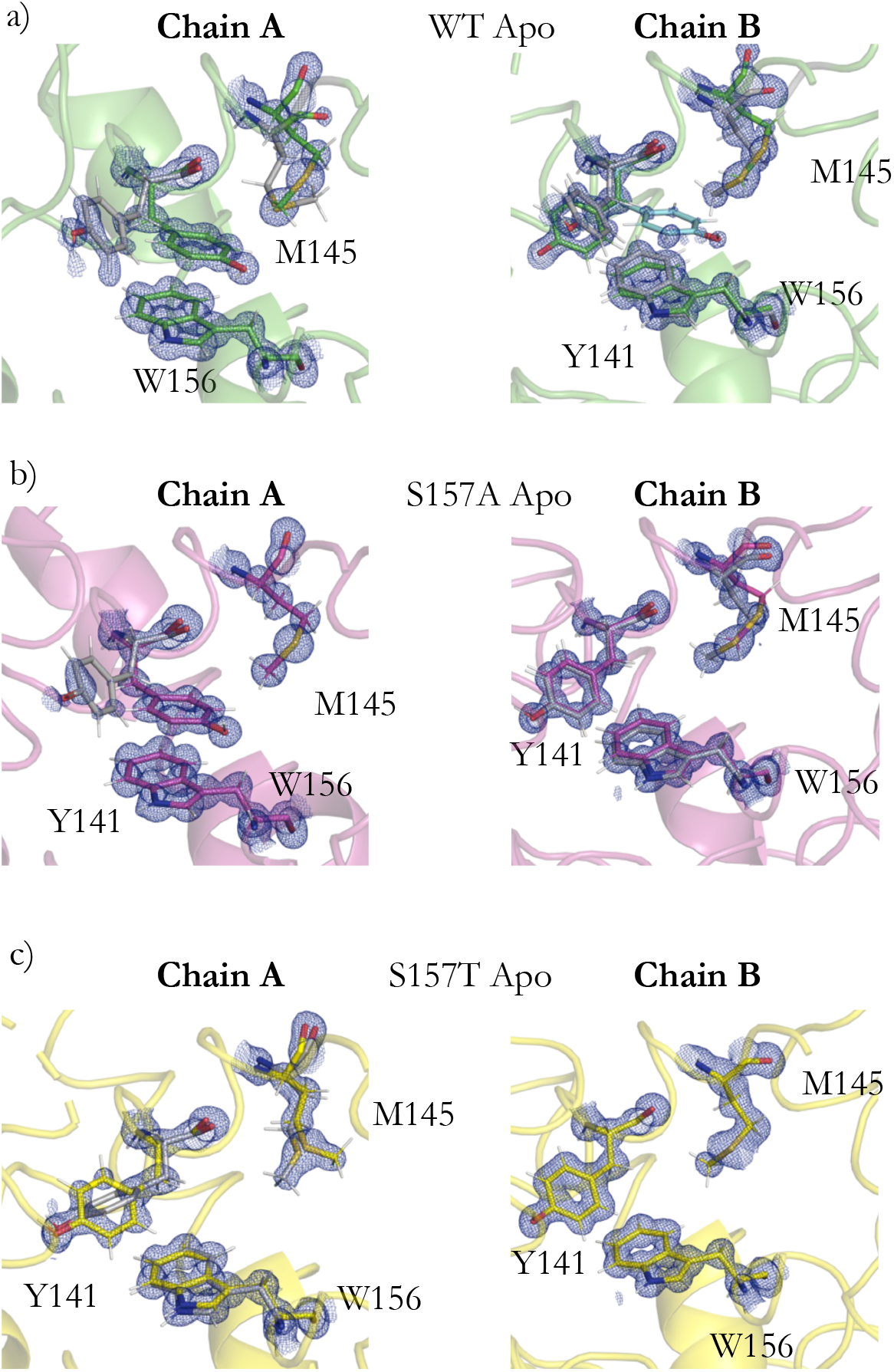
Conformations of residues at the dimer interface. Electron density (blue mesh) 2mF_obs_ – DF_calc_, contoured at 1 *σ*. Residues 148-153 are omitted for clarity. a) WT Apo state shows Tyr141 and Met145 can be found in multiple conformations, the minor state is coloured in grey. Chain A major conformation has Tyr141 found in the substrate primed conformation, whereas in chain B Tyr141 is only found in this conformation in a very low occupancy state. b) Ser157Ala has two conformations for Tyr141, the major π-stacking with Trp156, consistent with substrate primed state, and one conformation of Met145 in chain A, whereas there are two conformations in Chain B for Met145 and Try141 it does not π-stack with Trp156. c) Ser157Thr shows the side chain of Met145 is completely different to the WT and Ser157Ala mutant, neither chain shows Tyr141 in the substrate primed conformation.

To further investigate the impact of Ser157 mutations on the catalytic activity, crystals of both variants were soaked with ClAc for varying durations (between 10 seconds, and 48 hours). Short soaks of Ser157Ala with ClAc revealed substrate bound in the active site, positioned identically to that observed in the WT. However, longer soaking of both WT and Ser157Ala resulted in a loss of crystallinity, and high-quality diffraction data could not be obtained. In contrast, after a 48-hour soak of Ser157Thr crystals revealed formation of a covalent intermediate (Figure 5). This intermediate resembles the covalent intermediate previously observed in both active sites of the His280Asn variant, where lack of the active site base stalls the reaction at this point, and does not allow product to be released. Similarly, in Ser157Thr, the covalent intermediate forms in both chains, corroborating the previous work that the enzyme begins catalysis in the second chain before product release in the first chain. This highlights the significance of residues at the dimer interface modulating enzyme activity and confirming their role in the allosteric communication pathway between the two protomers. Similarly, in Ser157Thr, the covalent intermediate forms in both chains. Together, the assay and structural data indicate that Ser157Ala is proceeding through its reaction coordinate as well as speeding up the reaction relative to the WT. Ser157Thr can bind substrate and initiate the reaction, in contrast, it fails to release the covalent intermediate, preventing product (glycolate) formation, and demonstrates a new distinct conformation.

**Figure 5.**
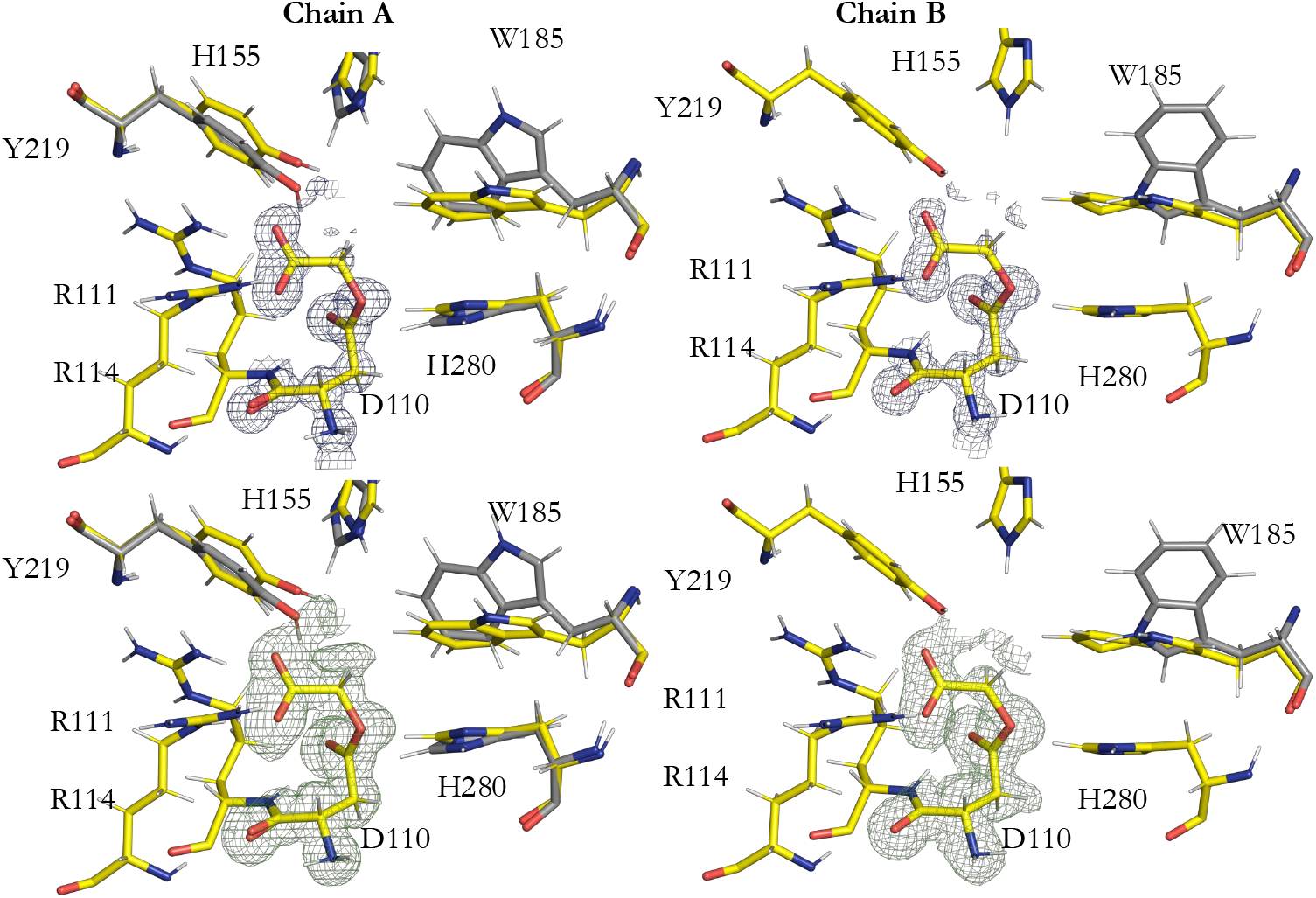
Electron density of the FAcD Ser157Thr covalent intermediate at 1.1 Å. Conformer A is in yellow, and Conformer B is in grey. Electron density (blue mesh) 2mF_obs_ – DF_calc_, contoured at 1.5 *σ*. Polder map of the complex (green mesh) at 3 *σ*.

Overall structural analysis shows that both mutations change the B-factor distributions of the refined models. As all datasets are of comparable resolution and quality, these differences are likely biologically relevant rather than artifacts of data collection. In WT FAcD, the average B-factor is higher for the B-chain than the A-chain. This chain-specific difference is also observed in the Ser157Ala and Ser157Thr variants, though it is less pronounced (Figure 6). Structurally, the Ser157Thr mutant adopts a conformation distinct from both WT and Ser157Ala, with a main chain offset of approximately 1 Å.

**Figure 6.**
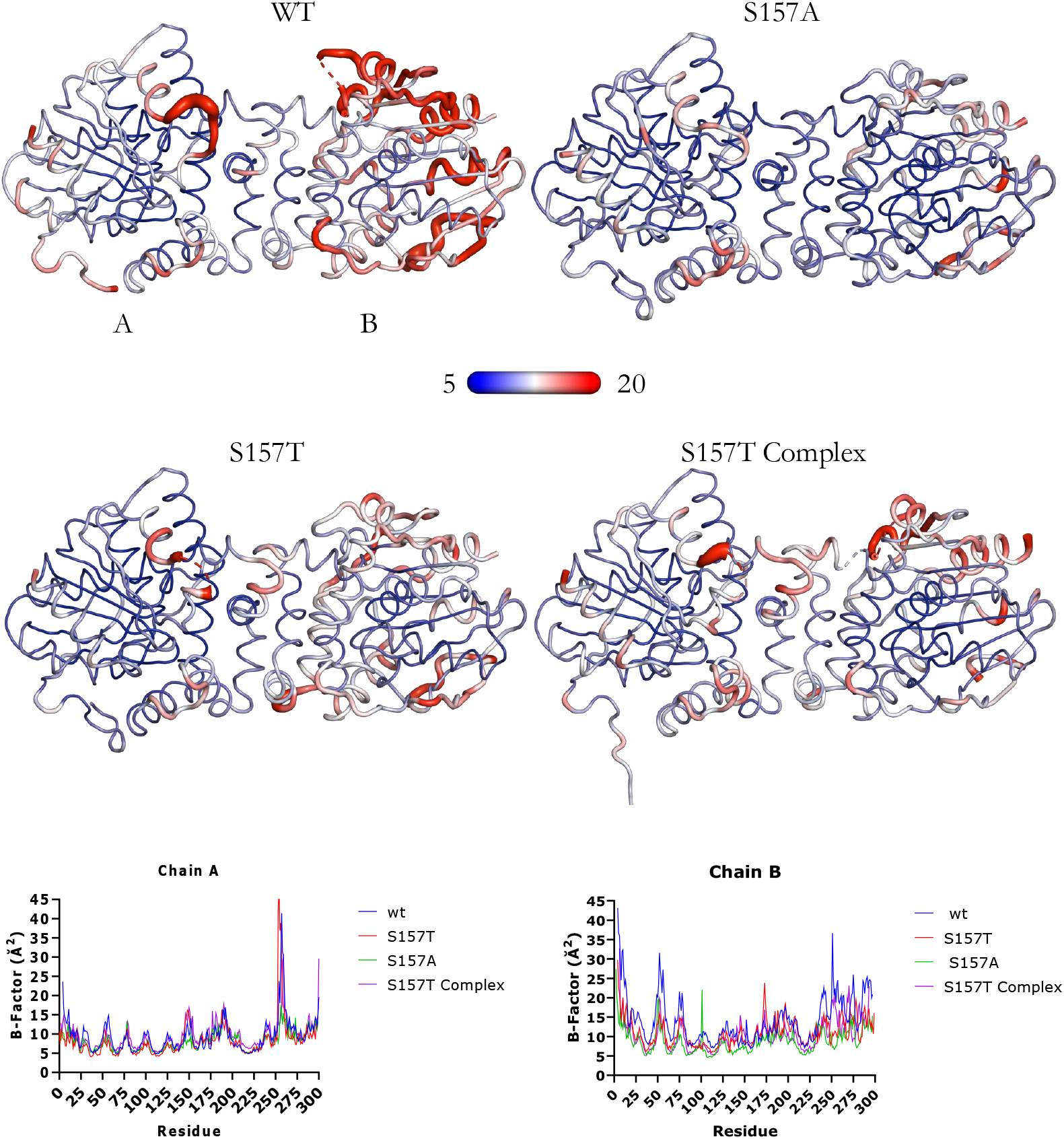
Overall structural analysis of FAcD and mutants. Crystallographic B-factors are coloured from blue (5) to red (20) with a putty representation and plotted as a function of residue. B-factors are noticeably different in the A and B as observed previously, but are also different lower for both mutants than wt.

## Discussion

As demonstrated by the kinetic assays, removal of the hydroxyl group from the Ser157 side chain increases the maximal rate of the reaction. This finding provides insight into how enzyme efficiency can be enhanced, particularly in the context of optimizing FAcD for bioremediation. In contrast, the Ser157Thr mutation, which retains the hydroxyl group but restricts side chain motion, curtails enzyme turnover. These results highlight the critical role of Ser157 in catalysis – preserving the chemical functionality but limiting its range of motion disrupts activity, while eliminating the functional group altogether enhances of the reaction rate. This highlights that FAcD has not evolved to turnover substrate as effectively as possible, unsurprising given the toxic nature of the substrate and the production of acid as one of the products. It is unlikely that FAc and ClAc are the natural substrates.

While the water coordination for the Ser157Ala mutant remains similar to WT, in Ser157Thr, the interface waters are suboptimally coordinated. Upon first inspection, these waters appear to have reduced occupancy and are displaced relative to their positions in WT. Closer inspection shows that W1 is completely absent, and W2 becomes more dynamic, adopting two distinct conformations. Another consequence of the Ser157Thr mutation is the loss of the minor conformation of Tyr141 that coordinates W1. Tyr141 has previously been shown to shift position depending on the reaction state, with molecular dynamics (MD) simulations indicating that its movement drives conformational selection ^11^. When Tyr141 adopts this minor conformation and π-stacks with Trp156, FAcD is primed for substrate binding ^3^. Despite this, crystal structures of Ser157Thr soaked with chloroacetate for 48 hours are capable of forming the covalent intermediate, that His280Asn is also capable of, showing that the mutant can still bind substrate and initiate catalysis, but it is unable to release the product. This suggests that Ser157Thr mutant can still reposition the Tyr141 side chain to sample the ligand-bound state, but this conformation is not predominantly sampled.

Mutations at the dimer interface alter FAcD’s ability to communicate across the subunits, a change that is evident in the structural data. Water bridges between the interface, particularly between the S157 side chains, are clearly observed, and the significance of these water networks was corroborated in MD simulations ^11^. The importance of hydrogen-bonding networks in mediating communication across oligomers has been demonstrated in other systems ^12^; for example, work on human transketolase identifies that low-energy hydrogen bonds and proton wires are capable of coupling two active sites together despite large-distances between them. While the water network in FAcD is obviously important, these findings also raise questions about the specific role of Ser157’s chemical functionality. Notably, the presence of a hydroxyl group is not strictly necessary for the water network to exist and to transmit information, as seen in the Ser157Ala mutant. However, reducing the mobility of this side chain, as seen in the Ser157Thr mutant, severely impairs enzyme activity. This suggests that the hydroxyl group may play a regulatory role. Is the role of this hydroxyl side chain to slow down the reaction? In systems where water wires are essential for communication, the chemical functionality of such residues is often crucial, indicating that Ser157 may serve a similar purpose in FAcD. Although this idea would predict the activity of the Ser157 mutants to be reversed, with Ser157Ala being inactive, where Ser157thr would maintain or enhance enzyme turnover. Clearly more understanding of how allosteric information is communicated is required to fully understand enzyme catalysis.

The Ser157Ala mutant is more efficient than WT and appears to adopt a single conformation at the dimer interface, whereas the WT is found in two conformations. This raises the question: could the Ser157Ala mutant be preferentially sampling the active conformation, whilst the WT samples an inactive and active conformation? In contrast, the Ser157Thr mutant adopts a substantially different conformation at the dimer interface that does not align with either WT or the Ser157Ala mutant. This structural difference may explain why Ser157Thr mutant is incapable of completing the reaction, as it is unable to sample the active conformation(s) that are predominant in the Ser157Ala and WT.

Crystallographically, it is not possible to distinguish whether the observed Ser157Thr structure represents a very slow mutant or if the reaction is stalling at the final step. However, based on kinetic data, the Ser157Thr mutant appears completely catalytically inactive. From this, we conclude that Ser157Thr behaves similarly to the His280Asn mutant, where the reaction stalls at the last step of hydrolysis as the active site base, His280, is unavailable to activate the catalytic water in the active site. In the Ser157Thr variant the position of the His280 side chain is unchanged, thus how the Ser157Thr mutation prevent the active site catalytic water from becoming catalytically active currently remains unclear. As both active sites are populated with the covalent intermediate, this could suggest that communication between the active sites is maintained. If the mutation has removed communication entirely, only one chain would be populated with the covalent intermediate. However, as both chains are capable of forming the covalent intermediate, this suggests that the first active site is capable of allosterically communicating with the second active site to initiate substrate turnover. Alternatively, this mutation could disrupt communication, meaning that both chains are capable of initiating the reaction independently of each other.

This work demonstrates the roles of residues at the protein interface and the regulation of enzyme catalysis by this interface interactions. Although our understanding of how these mutations impact enzyme catalysis is still unclear.

## Methods

### Protein purification

The mutations Ser157Thr and Ser157Ala were separately introduced into FAcD WT (modified pET 11) using site directed mutagenesis and the resulting variant proteins were expressed in BL21 DE3, purified via His-affinity, and size-exclusion chromatography in a final buffer of 50 mM TRIS-HCl pH 8.5, 150 mM NaCl, and crystallised as previously described ^1^.

### Data collection and model building

300 µm FAcD WT, Ser157Thr and Ser157Ala crystals were soaked in mother liquor supplemented with 30% *v/v* PEG400 with or without chloroacetate. Crystals were flash cooled in LN_2_ and diffraction data were measured at 100 K on beamline P14, EMBL@PetraIII. Data were collected using the Global Phasing Pipeline, with a beam energy of 18 keV on a EIGER2 X CdTe 16M detector ^13,14^. Data were processed with AutoPROC and STARANISO, with iterative model building and refinement with REFMAC5, and BUSTER ^10,15,16^.

### Enzyme activity

Absorbance measurements were performed using a Tecan Spark in the Protein production and characterisation facility (PPCF) at the CSSB Hamburg, Germany. Assays were conducted at 37 °C with 20 µgml^-1^ phenol red with varying concentrations of chloroacetate to a final volume of 200 µL in 1 mM TRIS, HCl, pH 8.5 buffer. FAcD was injected into the reaction to a final concentration of 5 µM (in 10 mM TRIS HCl pH 8.5), and the equivalent volume of 10 mM TRIS HCl pH8.5 was injected into the control reactions. The activity of FAcD was monitored by following the decrease in absorbance at 570 nm, corresponding to the change in pH ^1^. Proton concentration was calculated using the Henderson-Hasselbach equation^17^. Initial rates were calculated as number of protons produced in the first 1020 s of the reaction, and the rate of the control reaction was subtracted from the corresponding reaction to give a corrected initial rate. GraphPad prism was used with “substrate inhibition” equation, and parameters were calculated to fit a Michaelis-Menten function with a cut-off of 100 mM chloroacetate.

